# CRISPR/Cpf1 mediated genome editing enhances *Bombyx mori* resistance to BmNPV

**DOI:** 10.1101/2020.03.23.003368

**Authors:** Zhanqi Dong, Qi Qin, Zhigang Hu, Xinling Zhang, Jianghao Miao, Liang Huang, Peng Chen, Cheng Lu, Minhui Pan

## Abstract

CRISPR/Cas12a (Cpf1) is a single RNA-guided endonuclease that provides new opportunities for targeted genome engineering through the CRISPR/Cas9 system. Only AsCpf1 have been developed for insect genome editing, and the novel Cas12a orthologs nucleases and editing efficiency require more study in insect. We compared three Cas12a orthologs nucleases, AsCpf1, FnCpf1, and LbCpf1, for their editing efficiencies and antiviral abilities *in vitro*. The three Cpf1 efficiently edited the BmNPV genome and inhibited BmNPV replication in BmN-SWU1 cells. The antiviral ability of the FnCpf1 system was more efficient than the SpCas9 system after infection by BmNPV. We created FnCpf1×gIE1 and SpCas9×sgIE1 transgenic hybrid lines and evaluated the gene editing efficiency of different systems at the same target site. We improved the antiviral ability using the FnCpf1 system in transgenic silkworm. This study demonstrated use of the CRISPR/Cpf1 system to achieve high editing efficiencies in the silkworm, and illustrates the use of this technology for increasing disease resistance.

**Author Summary:** Genome editing is a powerful tool that has been widely used in gene function, gene therapy, pest control, and disease-resistant engineering in most parts of pathogens research. Since the establishment of CRISPR/Cas9, powerful strategies for antiviral therapy of transgenic silkworm have emerged. Nevertheless, there is still room to expand the scope of genome editing tool for further application to improve antiviral research. Here, we demonstrate that three Cpf1 endonuclease can be used efficiency editing BmNPV genome *in vitro* and *in vivo* for the first time. More importantly, this Cpf1 system could improve the resistance of transgenic silkworms to BmNPV compare with Cas9 system, and no significant cocoons difference was observed between transgenic lines infected with BmNPV and control. These broaden the range of application of CRISPR for novel genome editing methods in silkworm and also enable sheds light on antiviral therapy.

## Introduction

Genome editing introduces DNA mutations in the form of insertions, deletions or base substitutions within selected DNA sequences [1]. Clustered regularly interspaced short palindromic repeats (CRISPR) gene editing technology has been used in gene function research, genetic improvement, modelling biology and gene therapy [2-5]. Three effector proteins of class 2 type V CRISPR systems, the CRISPR/CRISPR-associated 12a (Cas12a, known as Cpf1) proteins of *Lachnosperaceae bacterium* (LbCpf1), *Francisella novicida* (FnCpf1) and *Acidaminoccocus sp*. (AsCpf1), have been shown to efficiently edit mammalian cell genomes with more efficient genome editing than the widely used *Streptococcus pyogenes* Cas9 (SpCas9) [6-8]. However, the CRISPR/Cpf1 system was rarely used for insect genome editing research and antiviral therapy [9].

The silk industry (*Bombyx mori, B. mori*), suffers great economic losses due to *B. mori* nucleopolyhedrovirus (BmNPV) infection[10-12]. CRISPR genome editing is an efficient and widely used technology for anti-BmNPV gene therapy, viral gene function research, and screening of potential targets in BmNPV infection[11, 13]. We first reported on highly efficient virus-inducible gene editing system, which demonstrated that CRISPR/Cas9 could edit the BmNPV genome and effectively inhibit virus proliferation[14]. Chen *et*.*al* effectively inhibited BmNPV proliferation and replication by editing the *ie-1* and *me53* of BmNPV immediate early genes in transgenic silkworm[15]. We improved the antiviral ability of transgenic silkworm by nearly 1,000 -fold by editing the two target sites of the *ie-1* gene to produce a large fragment deletion [11]. The CRISPR/Cas9 gene editing technology has also been used in antiviral resistance breeding by editing host factor and viral key genes in BmNPV infection. However, the antiviral resistance level using this system has now reached a plateau [13, 16, 17].

The CRISPR-Cas12a system (Cpf1) is a single RNA-guided endonuclease used for genome editing[6]. The Cpf1 enzyme has several gene editing characteristics that differ from the Cas9 system[7, 18]. One major difference between the Cpf1 and Cas9 systems is that Cpf1 recognizes a T-rich protospacer-adjacent motif (PAM), while Cas9 recognizes a G-rich PAM [19]. The Cpf1 system increases the potential targets sites that can be used for CRISPR-mediated gene editing[19]. Cpf1 enzyme requires one U6 (Pol-III) promoter to drive small CRISPR-derived RNA (crRNAs, 42-44-nt per-crRNA, 19-nt repeat and 23–25-nt spacer). However, the crRNA of the Cas9 enzyme requires an additional trans-activating crRNA (tracrRNA) to form the guide RNA[2, 19].Multiple crRNAs can be expressed as a single transcript to generate functional individual crRNAs after processing through Cpf1 nuclease. This can increase the efficiency of crRNA entry into cells[6, 20]. Cpf1 nuclease also generates a 5-bp staggered DNA double-strand break ends that are formed downstream of the PAM sequence, while Cas9 nuclease only formed a blunt-end cut 3 bp upstream of the PAM sequence[20, 21]. The unique editing features of the Cpf1 system, are conducive to overcoming the limitations of the Cas9 system.

We investigated the ability of AsCpf1, FnCpf1 and LbCpf1 to edit BmNPV genomes in B. mori. Our goals were to compare the Cpf1 and Cas9 systems for gene editing efficiency in anti-BmNPV therapy and to develop transgenic silkworms with BmNPV resistance. Initially, an AsCpf1, FnCpf1 and LbCpf1-based gene editing vector and the crRNA expression cassette were developed. Then, different Cpf1 nuclease activity with crRNA derived by the U6 promoter was evaluated for gene editing efficiency and antiviral ability in vitro. The antiviral abilities of FnCpf1 and SpCas9 systems, which are widely used for BmNPV genome editing, were compared. Finally, the gene editing efficiency and resistance level of the transgenic FnCpf1 and SpCas9 lines were evaluated by mortality analyses, sequencing and viral gene transcription in transgenic silkworms.

## Results

### CRISPR/Cpf1 system enables edit BmNPV genome

To determine whether the CRISPR/Cpf1 system could be used for gene editing in silkworm, we examined the functionality of three Cpf1 enzymes, AsCpf1, FnCpf1 and LbCpf1, which have been used to edit the genomes of mammal cells. We constructed a AsCpf1, FnCpf1 and LbCpf1 expression cassette attached to nuclear localization signal (NSL) and 3×HA tag, which gene is driven by the OpIE2 promoter and terminator by the OpIE2-PA. The crRNA expression cassettes consisting of a 20-21-nt direct repeat and a 23-nt guide sequence were arranged in tandem and driven by a signal U6 promoter of *B. mori*. Then, we transfected BmN-SWU1 cells with individual Cpf1 orthologs and gRNA to target endogenous loci containing the 5′ T-rich PAMs (Fig 1A).

**Fig. 1.**
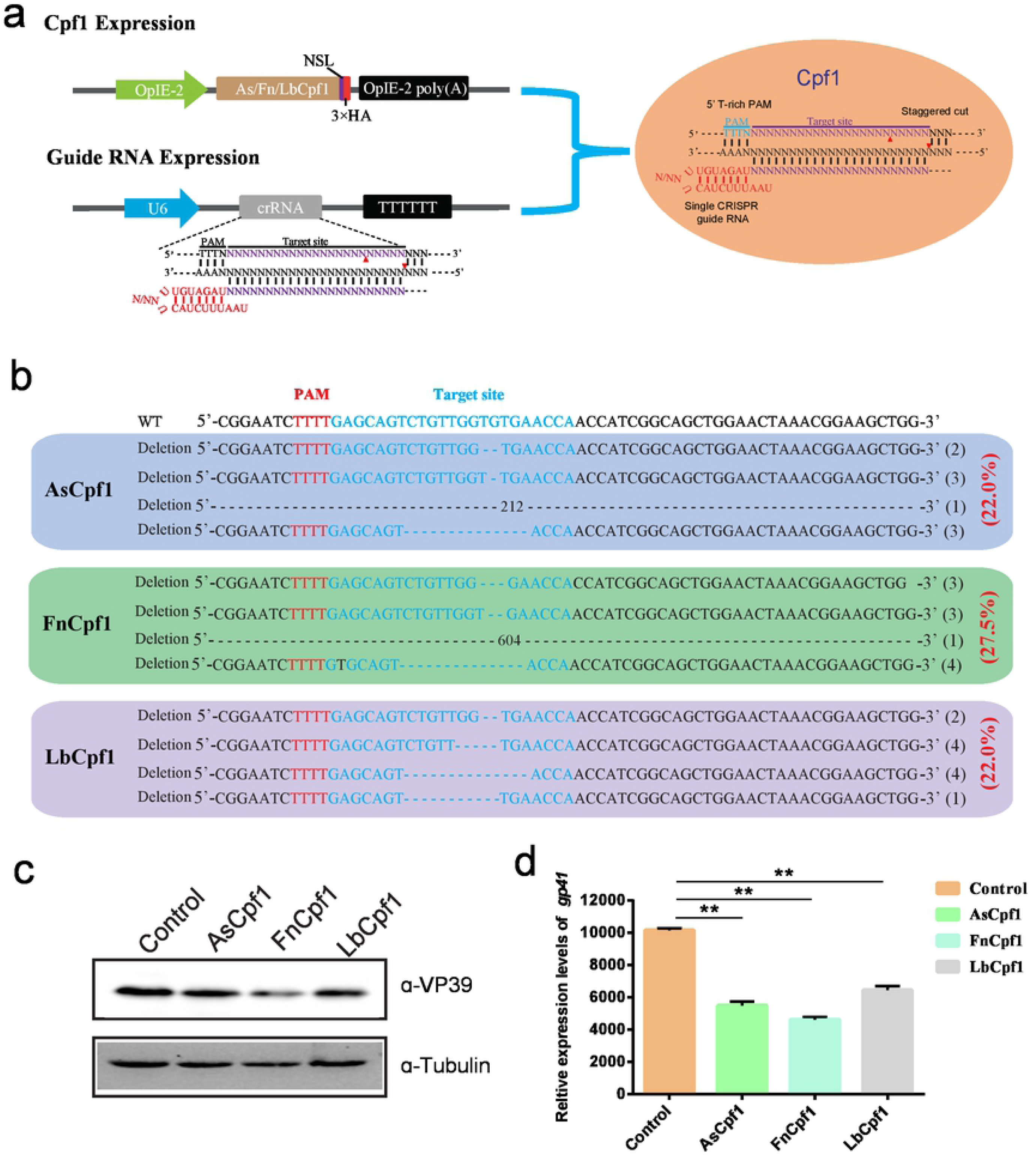
CRISPR/Cpf1 system enables editing BmNPV genome. (A) Cpf1 expression Cassette and crRNA target gene site. (B) DNA sequencing analysis of high-frequency genome mutations by Cpf1 in BmN-SWU1 cells. The BmNPV *ie-1* gene sequence is shown in black on top, the target site of gRNA is in blue, PAM sequence is in red, and the deletion sequence is indicated by dashes. (C) Western blot analysis of CRISPR/Cpf1 system mediated antiviral activity was monitored by the levels of the VP39 (top) and Tubulin (bottom). (D) DNA replication analysis of CRISPR/Cpf1 system mediated antiviral activity was monitored by the copies of gp41. Error bars represent standard deviations of three biological replicates. ** represents a statistically significant difference at *P*< 0.01.

Editing the BmNPV *ie-1* gene can effectively inhibit viral replication. We selected the *ie-1* gene as a target for further analysis. To facilitate the quantification and comparison of these nucleases, we constructed one vector system containing Cpf1 orthologs and gIE1. After transfecting the Cpf1 system in BmN-SWU1 cells, Sanger sequencing analysis revealed that all of the Cpf1 enzymes could edit the target site of the *ie-1* gene. The AsCpf1, FnCpf1 and LbCpf1 systems gene editing efficacy of the putative cleavage site reached 22.0%, 27.5% and 22.0% (Fig 1B). To further analyze whether the Cpf1 system could inhibit virus proliferation, we examined the change in VP39 protein expression after BmNPV infection. Western blot results showed that the expression of VP39 protein was significantly affected by eliminating the viral genome at 48 h p.i.. The VP39 protein expression levels were equivalent to 74.0%, 56.0% and 61.0% of the control, respectively (Fig 1C). To demonstrate the antiviral efficiency of the CRISPR/Cpf1 system, we also determined the replication of the viral genome through qPCR analysis. The amount of BmNPV DNA was affected after eliminating the viral genome. Compared with the control group, the AsCpf1, FnCpf1 and LbCpf1 systems was reduced BmNPV DNA by 46.0%, 54.6% and 36.5%, respectively (Fig 1D). All of the three constructed CRISPR/Cpf1 gene editing systems significantly inhibited virus replication in *B. mori*, and the FnCpf1 system had the greatest antiviral effect.

### Analysis of antiviral ability of CRISPR/Cas9 and CRISPR/Cpf1 in vitro

To evaluate the performance of the different CRISPR systems in *B. mori*, we focused on FnCpf1 and SpCas9 gene editing systems for the same target site. We initially chose the BmNPV *ie-1* gene as the target. PAM profiling of FnCpf1 and SpCas9 is shown in Fig 2A. After transfected with FnCpf1×gIE1 and SpCas9×sgIE1 in BmN-SWU1 cells, the cells infected with the vA4^prm^-EGFP virus at MOI of 10. At 48 h p.i., viral DNA replication showed that different gene editing systems could significantly inhibit BmNPV DNA replication, and the FnCpf1 system had a greater inhibition effect compared with the SpCas9 system (Fig 2B). BmNPV DNA replication levels were reduced by 54.6% in the FnCpf1 system relative to the control and decreased more than 38.5% compared with the SpCas9 system. We analyzed the changes of VP39 protein expression in the FnCpf1 and SpCas9 systems. The Western blot analysis showed that the FnCpf1 and SpCas9 system could significantly inhibit VP39 protein expression. The FnCpf1 system only detected the VP39 protein at 48 h p.i. (Fig 2C). After the FnCpf1 system was transfected in BmN-SWU1 cells and infected with BmNPV, no significant VP39 protein expression was detected in the FnCpf1 system at 0-24 h p.i.; however, a weak VP39 protein band was able to detected in the SpCas9 system at 24 h p.i.. The VP39 protein expression of the FnCpf1 system was also lower than that of SpCas9 system at 48 h p.i. (Fig 2C). These results demonstrated that the antiviral ability of the FnCpf1 system was more effective than the SpCas9 system for BmNPV at the same target site.

**Fig. 2.**
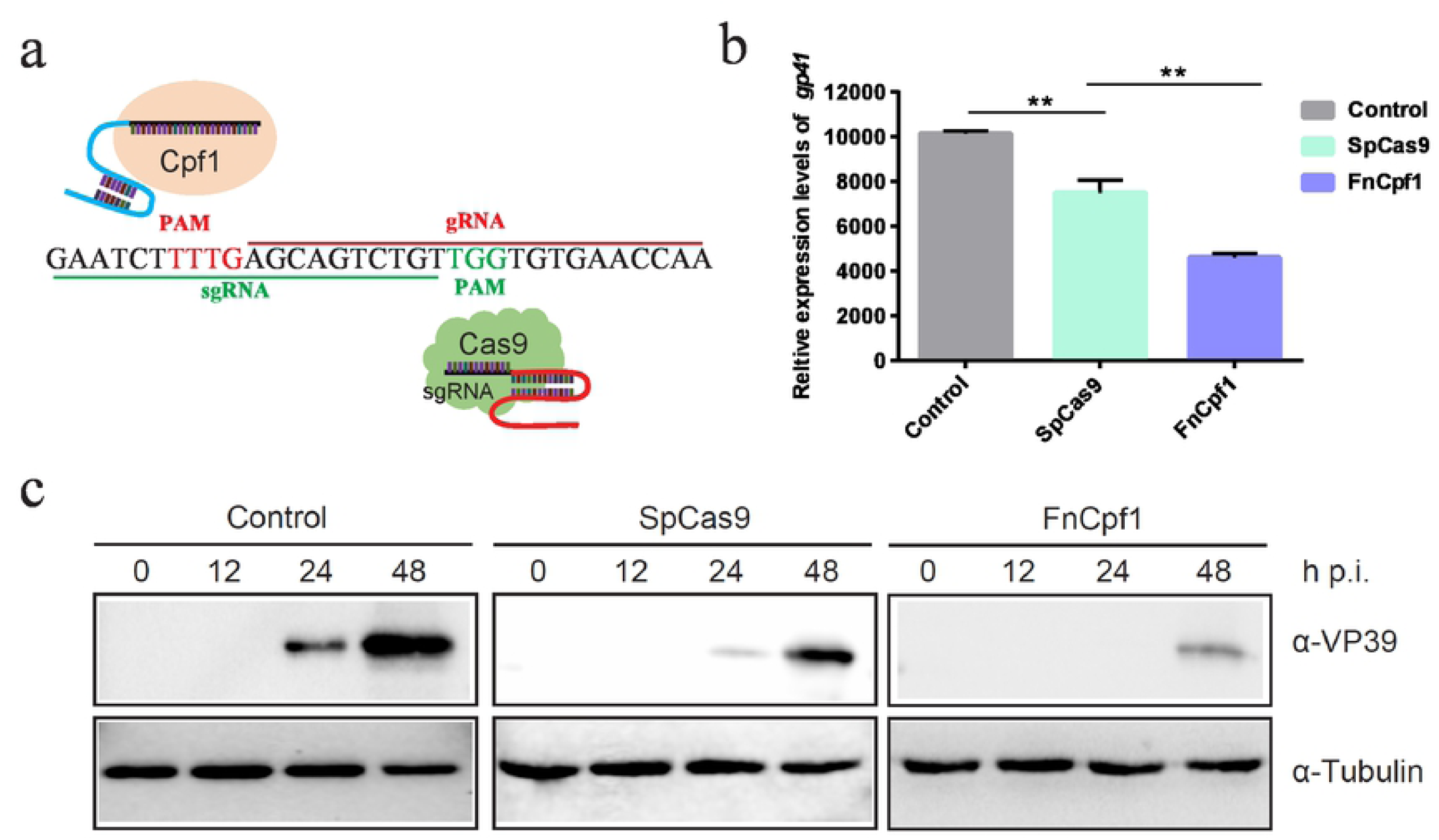
Comparative analysis of the antiviral ability of CRISPR/Cas9 and CRISPR/Cpf1 *in vitro*. (A) Comparison of CRISPR-Cas9 versus CRISPR-Cpf1 mediated genome editing. Cpf1, Cas9, crRNA and PAM are shown. (B) Analysis of BmNPV DNA replication in CRISPR/Cpf1 and CRISPR/Cas9 system. Error bars represent standard deviations of three biological replicates. ** represents a statistically significant differences at *P*< 0.01. (C) Western blot analysis of CRISPR/Cpf1 and CRISPR/Cas9 system mediated antiviral activity was monitored by the levels of the VP39 (top) and Tubulin (bottom). The ratios of different types of mutations.

### Gene editing efficiency of CRISPR/Cas9 and CRISPR/Cpf1 systems in transgenic silkworm

To compare genome editing efficiency of CRISPR/Cpf1 and CRISPR/Cas9, we constructed FnCpf1 and SpCas9 system transgenic vectors. The vectors pBac[IE2-FnCpf1-OPIE2-PA-3×P3 EGFP afm], pBac[U6-gIE1-3×P3 DsRed afm], and pBac[U6-sgIE1-3×P3 DsRed afm] expressed the FnCpf1 protein, gRNA and the sgRNA target sequence, respectively. The SpCas9 transgenic line was studied previously [11].

After selection of the FnCpf1, gIE1, SpCas9 and sgIE1-positive transgenic lines, double-positive transgenic FnCpf1 × gIE1 and SpCas9 × sgIE1 lines were obtained through FnCpf1 and gIE1 or Cas9 and sgIE1 transgenic line hybridization (Fig 3A). The FnCpf1 × gIE1 line expressed both FnCpf1 protein and gIE1 target sequence, and the SpCas9 × sgIE1 line expressed both SpCas9 protein and sgIE1 target sequence. In the G2 generation, silkworms with both red fluorescent protein and green fluorescent protein expression in their eyes were the double-positive transgenic FnCpf1 × gIE1 or SpCas9 × sgIE1 lines (Fig 3A).

To compare the gene editing efficiency of CRISPR/Cas9 and CRISPR/Cpf1 system in transgenic silkworm, we selected the *ie-1* gene of BmNPV as the target gene site. The two systems targeted the same site of *ie-1*. After infection with OBs under the same conditions, we determined the gene editing efficiency of the target sites in the transgenic hybrid line, FnCpf1 × gIE1 or SpCas9 × sgIE1. Sequencing of PCR fragments from these lines demonstrated that both the CRISPR/Cas9 and CRISPR/Cpf1 gene editing systems were able to edit the *ie-1* gene in the BmNPV genome (Fig 3B). We also found that the sequence of SpCas9 × sgIE1 lines was able to edit the target site within the BmNPV genome, which mainly appeared as the absence of 3–30 bp, and only one colony had large deletions in all sequencing (Fig 3B). In contrast, most clones of the transgenic FnCpf1 × gIE1 line showed large deletions, ranging from 500 to 1400 bp. More than 80% large deletions accounted for all sequencing (Fig 3B).

**Fig. 3.**
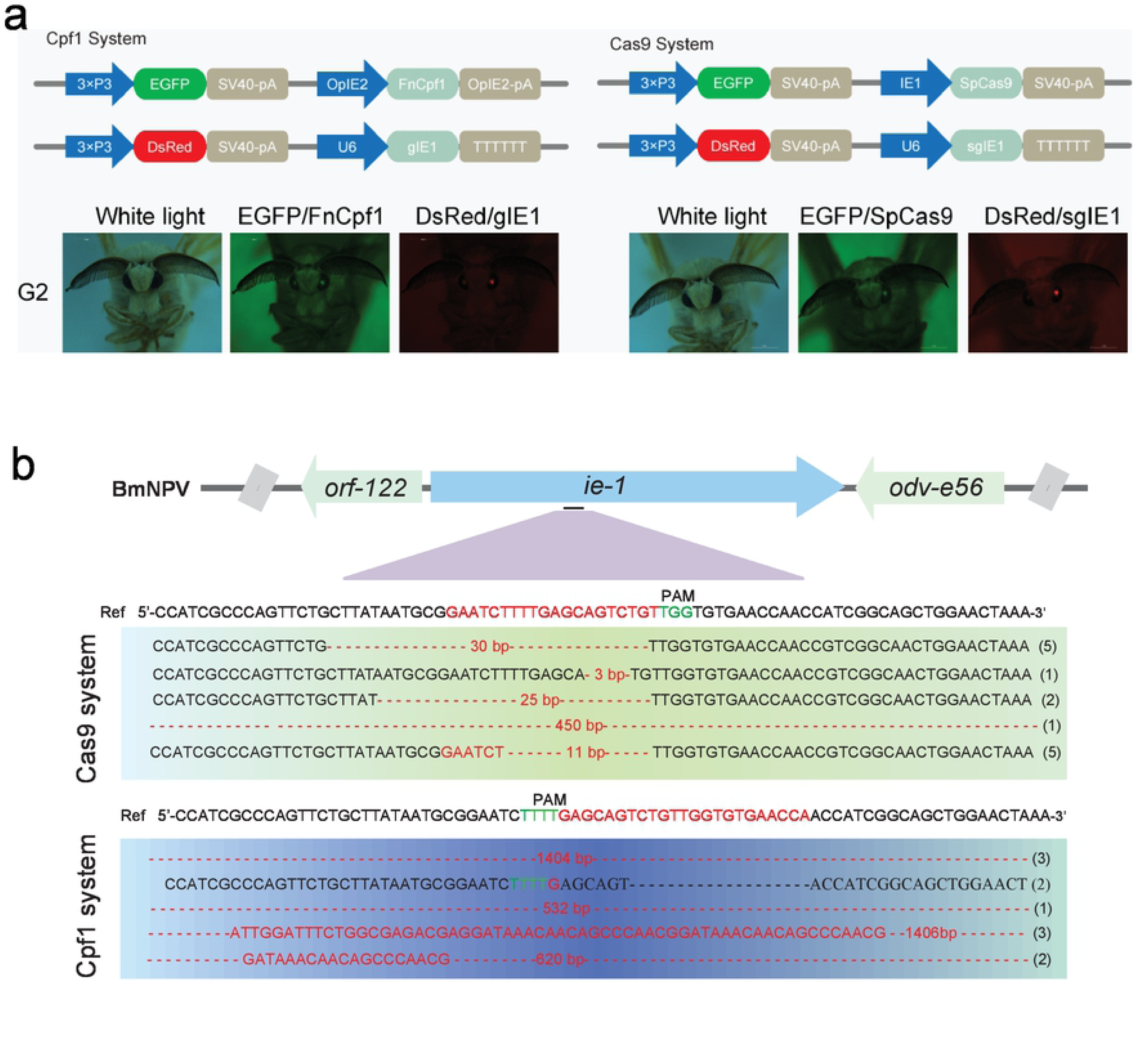
Comparison of gene editing efficiency of CRISPR/Cas9 and CRISPR/Cpf1 system in transgenic silkworm. (A) Schematic presentation of transgenic vector construction of pBac[OpIE2prm-Cpf1-OpIE2-PA-3×P3 EGFP afm], pBac[U6-gIE1-3×P3 DsRed afm] and pBac[U6-sgIE1-3×P3 DsRed afm] (top). Double positive individuals FnCpf1×gIE1 and SpCas9×sgIE1 obtained by were screened by fluorescence microscopy (bottom). (B) Sequencing results of two transgenic lines generated by mutagenesis at *ie-1* site.

To evaluate the potential off-target effects of FnCpf1, we examined all possible off-target sites with high sequence similarity to gIE1 in the silkworm genomes. We selected three non-specific editing sites with the highest similarity for further confirmation by PCR in transgenic lines. Among the three predicted off-targeting sites, we did not detect any off-target mutations in the FnCpf1 × gIE1 transgenic lines (Table 1). These results showed that the FnCpf1 systems used in antiviral research had no significant effects on non-specific loci even for editing a highly similar site in the silkworm.

### Silkworm resistance to BmNPV conferred by the CRISPR/Cpf1 system

We determined whether the FnCpf1 system could enhance antiviral activity compared with SpCas9 system in transgenic lines. The transgenic hybrid lines FnCpf1 × gIE1, SpCas9 × sgIE1 and DaZao were infected with 1 × 10^6^ OBs/larva by inoculating 4th instar larvae. Under these conditions, the FnCpf1 × gIE1 and SpCas9 × sgIE1 lines significantly reduced the BmNPV infection. The survival rate of the SpCas9 × sgIE1 lines was 59% until 10 d p. i., whereas the control had large-scale mortality after 5 to 10 d p. i. (Fig 4A). The survival rate of the FnCpf1×gIE1 lines were further increased when they were inoculated with OBs. The FnCpf1×gIE1 lines started to die on 6 d p. i., but the survival rate of the FnCpf1×gIE1 lines was still >65% after 10 d p. i. (Fig 4A). These results suggested that the CRISPR/Cpf1 system, in transgenic silkworm, could more effectively improve the antiviral activity (Fig 4A). We determined if the surviving transgenic FnCpf1 × gIE1 and SpCas9 × sgIE1 silkworm lines had altered cocoon characteristics after BmNPV infection. We compared the transgenic lines to the control, and found that they were similar with differences ranging from 11% to 18% (Fig 4B).

**Fig. 4.**
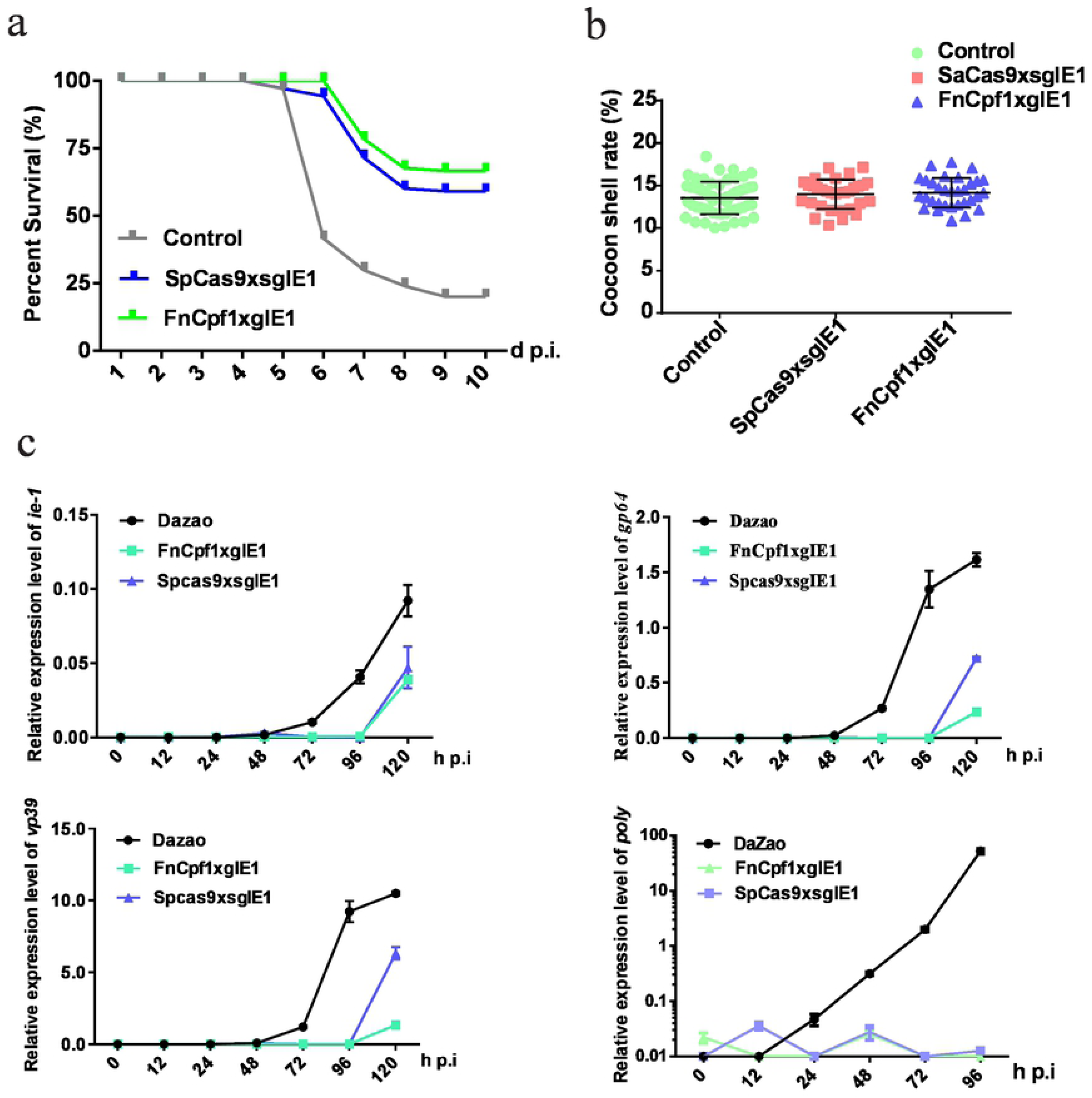
Improved silkworm resistance to virus conferred by CRISPR/Cpf1 system. (A) Survival rate of transgenic hybrid FnCpf1×gIE1 and SpCas9×sgIE1 lines after inoculation with 1 × 10^6^ OBs per 4^th^ instar larva. Each group included 30 larvae included the standard deviations of three biological replicates. The mortality was scored at 10 d p.i.. (B) Cocoon shell rate analysis of the FnCpf1×gIE1 and SpCas9×sgIE1 lines. Each value represents three biology replicates. NS, not significant. (C) Gene expression levels of *ie-1, gp64*, vp39 and *poly* after OB inoculation were analyzed by RT-PCR in two transgenic lines. The data represent of three experiments. NS, not significant. ** represents statistically significant differences at the level of *P*< 0.01.

To compare the antiviral ability of different gene editing systems, we also analyzed the changes of BmNPV gene expression levels at different stages. Similarly, DaZao, FnCpf1 × gIE1 and SpCas9 × sgIE1 transgenic lines inoculating 4th instar larvae with 1 × 10^6^ OBs/larva. At 0, 12, 24, 48, 72, 96, and 120 h p. i., total RNA was isolated from each transgenic line and the samples were analyzed by RT-PCR. We studied immediate early gene *ie-1*, early gene *gp64*, late gene *vp39*, and very late gene *poly* of BmNPV to analyze the viral expression levels at different stages. The RT-PCR results showed that the expression of *ie-1, gp64, vp39*, and *poly* genes were maintained at a very low level in the FnCpf1 × gIE1 and SpCas9 × sgIE1 transgenic lines after BmNPV infection. However, the viral gene expression levels increased as expected in the control (Fig 4C). The viral gene expression levels of FnCpf1 × gIE1 and SpCas9 × sgIE1 transgenic lines were 10^4^–10^5^-fold lower compared with the DaZao lines. The FnCpf1 × gIE1 lines had a 10-fold reduction in viral expression compared with the SpCas9 × sgIE1 at different stages.

## Discussion

Genome editing has the potential to accurately edit the genomes of model organism [2, 20, 22]. Cas9, Cas12a (Cpf1), Cas12b, Cas13, Cas3 and Cas14 based CRISPR systems have been explored for editing human, animals, plants and microbe genomes[23-28]. Cpf1 is a type V CRISPR-effector protein with greater specificity for genome editing in mammals and plants[6, 7]. To overcome the limitation of Cas9 for antiviral research in *B. mori*, we engineered an improved CRISPR/Cpf1 system and used it to evaluate its efficiency and accuracy for BmNPV genome editing. This application expands the used of CRISPR technology in insect.

RNase activity of AsCpf1, FnCpf1 and LbCpf1 has been used for genome editing[19, 21, 29]. AsCpf1 previously known to efficiently edit insect genomes. It was also not known which Cpf1 system has higher editing efficiency in Lepidoptera species, such as *B. mori*. We compared the ability of AsCpf1, FnCpf1 and LbCpf1 to edit genomes in BmN-SWU1 cells. By determining the editing efficiency and antiviral ability, we showed that three Cpf1s can induce heritable mutations at target sites (Fig 1B). Large fragment deletions occurred in the PCR products of AsCpf1 and FnCpf1. FnCpf1 was the most efficient gene editing system studied (Fig 1B). To avoid the effect of the target site on gene editing efficiency and antiviral ability, we designed the target site of the Cas9 system to be the same site (Fig 2A). These findings emphasize that FnCpf1 system has potential for use in the development of virus-resistant silkworm lines.

Cas9 system transgenic positive lines can fully edit the target gene [11]. To determine the reason for the difference in antiviral abilities of the Cas9 and Cpf1 systems, we constructed transgenic lines. The transgenic FnCpf1×gIE1 lines could create larger fragment deletions compared with SpCas9×sgIE1 lines. Based on the gene-editing principles of the Cas9 and Cpf1 systems, we believe that the cleavage site was distant from the target site of Cpf1 system, and the target site was not destroyed after cleavage[6]. After target cleavage, it could produce double-strand breaks, which resulted in large fragments being deleted [21]. It also had a greater impact on the function of the viral gene, which could inhibit viral DNA replication. In contrast, Cas9 system produced blunt ends after editing, and was easily repaired by homologous recombination. The cleavage site of Cas9 was at the target site, resulting the system unable to recognize it again.

Silkworm selection for virus resistance is a traditional method used in the sericulture industry. Interfering with the key genes of BmNPV or overexpression of resistance genes can increase antiviral ability of the silkworm[10, 12, 30]. The CRISPR/Cas9 gene editing system had allowed us to improve the antiviral ability of transgenic silkworms. This is accomplished by editing the virus early transcriptional activators, multiple target sites and multiple genes editing, and editing host-dependent factors[13, 31]. Increased antiviral ability, using tradition means, has reached a limit, and new technology is needed to increase resistance to virus attack. We used three different Cpf1 systems for editing the BmNPV genome, and screened a Cpf1 system with high antiviral ability and gene editing efficiency in *B. mori*. This research demonstrated that the antiviral ability of Cpf1 system can be improved compared with the Cas9 system under the same target site in transgenic silkworms (Fig 4A). The Cpf1 system can drive many crRNAs through a U6 promoter. In further research we can edit the BmNPV genome through multiple genes and multiple target sites. This will increase the negative effects on the BmNPV genome and improve the virus resistance of transgenic silkworms. We can also try to edit multiple silkworm viruses by synthesizing more crRNAs (such as crRNAs of *B. mori* densovirus, *B*.*mori* cytoplasmic polyhedrosis virus and other infectious diseases of *B. mori*) to one vector. This will further expand the scope and efficiency of transgenic antiviral breeding.

In conclusion, we developed a novel CRISPR nuclease platform, AsCpf1, FnCpf1, and LbCpf1, which can be used for BmNPV genome editing and breeding of virus-resistant silkworms. Our research data indicated that the CRISPR/Cpf1 system is a powerful tool for silkworm selection. The system can be used to improve silkworm virus resistance and also as a way to combat other infectious diseases. The successful application of CRISPR/Cpf1 genome editing system can be used to address diseases in B. mori and perhaps other economically important insect.

## Methods

### Cells

A *B. mori* cell line BmN-SWU1, derived from ovary tissue, was maintained in our laboratory and used in this study [32]. BmN-SWU1 cell lines were cultured at 27°C in TC-100 medium (United States Biological, USA). The medium was supplemented with 10% (V/V) fetal bovine serum (FBS) (Gibco, USA).

### Viruses

A recombinant BmNPV (vA4^prm^-EGFP) was constructed and used in this study[33]. The baculovirus contained a gene encoding for an EGFP marker gene under the control of the *B. mori* actin A4 promoter. Budded virus (BV) amplification was performed by infection with BmN-SWU1 and harvested at 120 h post-infection (h p.i.). Viral titration was performed using the plaque assay method. Occlusion-derived virus (OB) amplification was performed using oral inoculation with the wild-type (WT) Chongqing strain of BmNPV in silkworm larvae. OBs were harvested from the infected hemolymph before the larvae died [34].

### Silkworm strains

The “DaZao” and transgenic Cas9 strain of *B. mori* were maintained in our laboratory[24]. Silkworm larvae were fed on fresh mulberry leaves and maintained at 25°C under standard conditions.

### Vector construction

To explore whether the CRISPR/Cpf1 system could be used for gene editing in *B. mori*, wild-type LbCpf1 plasmid, pY016 (pcDNA3.1-LbCpf1, Addgene plasmid # 69988), AsCpf1 plasmid, pY010 (pcDNA3.1-AsCpf1, Addgene plasmid # 69982) and FnCpf1 plasmid, pY004 (pcDNA3.1-FnCpf1, Addgene plasmid # 69976) were obtained from Addgene. AsCpf1, FnCpf1 and LbCpf1 fragment were cloned into pSL1180-IE2^prm^-OpIE2-PA vector by digested with *BamH I* and *Kpn I* restriction sites, yielding pSL1180-OpIE2^prm^-AsCpf1-OpIE2-PA, pSL1180-IE2^prm^-FnCpf1-OpIE2-PA and pSL1180-OpIE2^prm^-LbCpf1-OpIE2-PA. The crRNA expression cassette under the control of the *B. mori* U6 promoter was synthesized by BGI and named pSL1180-U6-gRNA. The candidate crRNA target sequences were designed using CRISPR design software https://crispr.cos.uni-heidelberg.de/index.html). We sequentially linked the U6-gRNA expression cassettes into the pSL1180-OpIE2^prm^-AsCpf1-OpIE2-PA, pSL1180-OpIE2^prm^-FnCpf1-OpIE2-PA and pSL1180-OpIE2^prm^-LbCpf1-OpIE2-PA, and then used restriction enzymes to verify cloning, respectively. Cas9 and sgRNA expression cassettes of the target gene *ie-1* used previous constructs. We selected the target sites of the BmNPV *ie-1* gene as CRISPR/Cpf1 and the CRISPR/Cas9 gene editing sites. Sequences for all of the targets of the guide RNAs are provided in S1 Table.

The transgenic silkworm Cas9 lines were constructed as previously reported. To obtain the green fluorescent protein transgenic vector pBac [OpIE2prm -FnCpf1-OpIE2-PA-3×P3 EGFP afm], the fragment OpIE2prm-FnCpf1-OPIE2-PA was ligated to the pBac [3×P3 EGFP afm] vector after a single digestion of pSL1180-OpIE2prm-FnCpf1-OpIE2-PA by *Asc I* restriction endonuclease. Simultaneously, *ie-1* target genes vector pSL1180-U6-gIE1 and pSL1180-U6-sgIE1 were ligated to a pBac [3×P3 DsRed afm] vector after single digestion with *Bgl II*, which generated a red fluorescent protein transgenic vector for pBac [U6-gIE1 DsRed afm] and pBac [U6-sgIE1-3×P3 DsRed afm]. All of the primers used are listed in S1 Table, and all of the constructed vectors were verified by sequencing.

### sgRNA and gRNA design

BmNPV *ie-1* genes were used as targets for gene editing. To avoid the influence of target sites on the editing efficiency of different gene editing systems, we chose the same site of *ie-1* (located at 360 transcription start site of *ie-1*) as the target site of the Cpf1 and Cas9 gene editing system. We predicted sgIE1 target gene sequences using an online analysis tool (http://crispr.dbcls.jp/) [35]. All of the candidate sgRNA target sequences have the GN19NGG sequence. The candidate gRNA target sequences were designed using a CRISPR design software tool (https://crispr.cos.uni-heidelberg.de/index.html)[36]. All of the candidate gRNA target sequences met the requirements of of TTTN PAM recognition domain.

### Quantitative PCR (qPCR) DNA replication assay

Total DNA was extracted from silkworm cells and larvae using a Wizard Genomic DNA extraction kit (Promega, USA). The copy number of BmNPV was calculated based on quantitative PCR as previously described [34]. PCR was performed in 15 μl reactions using 1 μl of extracted DNA solution as the template. All of the experiments were repeated three times.

### Western Blot analysis

After BmN-SWU1 cells were transfected with the indicated plasmids, the cellular protein was extracted in IP buffer containing 10 μl protease inhibitors (PMSF) and boiled for 10 min. Protein suspension samples were separated by 12% SDS-PAGE and then transferred to a nitrocellulose membrane. The membrane was incubated with mouse α-HA (1:2000; Abcam, UK), mouse α-PCNA (1:2000; Abcam, UK), rabbit α-Tubulin (1:5000; Sigma, USA) and rabbit α-VP39 (1:2000) for 1 h. Then, the membrane was further incubated with HRP-labeled goat anti-mouse IgG (1:20000; Beyotime, China) and HRP-labeled goat anti-rabbit IgG (1:20000; Beyotime) for 1 h. Finally, the signals on the membrane were visualized by Clarity Western ECL Substrate (Bio-Rad, USA) following manufacturer’s instructions. Tubulins were used to estimate the total protein levels.

### Mutagenesis analysis at target sites

The purified BmNPV genome DNA products were amplified by PCR, and the resulting products were ligated into a pEASY-T5 Zero cloning vector (TransGen Biotech, Beijing, China). The plasmid was analyzed by Sanger sequencing using M13 primers and aligned with the *ie-1* sequence. All of the primers used for detection are presented in S1 Table.

### Microinjection and screening

The transgenic vector pBac [OpIE2prm-FnCpf1-OpIE2-PA-3×P3 EGFP afm], pBac [U6-gIE1-3×P3 DsRed afm] and pBac [U6-sgIE1-3×P3 DsRed afm] were mixed with the helper plasmid pHA3PIG in the ratio of 1:1 and injected into silkworm eggs as previously described [11]. The positive individuals were screened by fluorescence microscopy. Double positive individuals FnCpf1×gIE1 were obtained by crossing FnCpf1 and gIE1, double positive individuals SpCas9×sgIE1 were obtained by crossing SpCas9 and sgIE1. All of the positive strains were identified by PCR amplification and fluorescence screening.

### Off-target assays

Potential off-target sites in the silkworm genome were predicted using CRISPR design software (http://crispr.dbcls.jp/)[35]. We screened three potential sites of gIE1 with the highest off-target efficiency and examined these by PCR amplification. The corresponding PCR products were sequenced, and then aligned with the IE1 sequence. All of the off-target site primers used in the study are presented in S1 Table.

### Mortality analyses

The OBs of BmNPV were purified from diseased larvae and stored at 4°C. The transgenic silkwormS FnCpf1×gIE1 and SpCas9×sgIE1 were inoculated with 1 × 10^6^ OBs/larva during the fourth instar. Each experimental group contained 30 larvae, and the test was performed in triplicate. Each experimental group was reared individually and we calculated the survival rate 10 d post-inoculation.

### Determination of expression levels by real time-PCR (RT-PCR)

Total RNA was isolated from each cell or leaves and the cDNAs were synthesized using a cDNA synthesis Kit (OMEGA, USA). Gene expression was determined by RT-PCR analysis using an Applied Biosystems 7500 Real-Time PCR System (Life Technologies, USA) with SYBR Select Master Mix Reagent (Bio-Rad). The housekeeping gene (*B. mori sw22934*) was used as a control. The normalized expression, reported as the fold change, was calculated for each sample using the 2^-ΔΔCT^ method. Three replicates were performed for each reaction. The RT-PCR specific primers are listed in S1 Table.

### Characteristics analysis of transgenic silkworm

The cocoon volumes of the two transgenic lines, FnCpf1×gIE1 and SpCas9×sgIE1 were analyzed after pupation. Each transgenic line, including 30 larvae, was characterized by the mean of three independent replicates. The cocoon shell rate was calculated as the combined pupa and cocoon weight.

### Statistical analysis

All of the data are expressed as mean ± SD of three independent experiments. Statistical analyses were performed with a two-sample Student’s *t-*test using GraphPad Prism 6. Differences were considered highly significant at *P* < 0.01.

## Acknowledgment

This work was supported by grants from the National Natural Science Foundation of China (Nos. 31902214 and 31872427), Fundamental Research Funds for the Central Universities (No. XDJK2020C010), Natural Science Foundation of Chongqing (cstc2019jcyj-msxm2371) and China Agriculture Research System (No. CARS-18).

## Author contributions

Z.D., Q.Q. and L.H. performed the vector cloning, sequencing, cell cultures and PCR. Z.D., Q.Q. and X.Z. performed the transgenic injection. J.M. and Z.H. performed the mortality analyses and DNA replication assay. Z.D., M.P. and C.L. conceived the experimental design and helped with data analysis. Z.D., M.P., P.C., and C.L. prepared of the manuscript. The final manuscript was reviewed and approved by all authors.

## Ethics approval and consent to participate

Not applicable.

## Supporting Information

**S1 Table Sequences of the primers used in this study**.

## References

1. Backes S, Hess S, Boos F, Woellhaf MW, Godel S, Jung M, et al. Tom70 enhances mitochondrial preprotein import efficiency by binding to internal targeting sequences. Journal of Cell Biology. 2018;217(4):1369-82. doi:10.1083/jcb.201708044. PubMed PMID: WOS:000428997800019.

2. Burgess DJ. Technology: a CRISPR genome-editing tool. Nat Rev Genet. 2013;14(2):80. doi:10.1038/nrg3409. PubMed PMID: 23322222.

3. Li F, Shi J, Lu HS, Zhang H. Functional Genomics and CRISPR Applied to Cardiovascular Research and Medicine. Arterioscler Thromb Vasc Biol. 2019;39(9):e188-e94. doi:10.1161/ATVBAHA.119.312579. PubMed PMID: 31433696; PubMed Central PMCID: PMCPMC6709691.

4. Alves-Bezerra M, Furey N, Johnson CG, Bissig KD. Using CRISPR/Cas9 to model human liver disease. JHEP Rep. 2019;1(5):392-402. doi:10.1016/j.jhepr.2019.09.002. PubMed PMID: 32039390; PubMed Central PMCID: PMCPMC7005665.

5. Beretta M, Mouquet H. [CRISPR-Cas9 editing of HIV-1 neutralizing human B cells]. Med Sci (Paris). 2019;35(12):993-6. doi:10.1051/medsci/2019196. PubMed PMID: 31903905.

6. Fagerlund RD, Staals RH, Fineran PC. The Cpf1 CRISPR-Cas protein expands genome-editing tools. Genome Biol. 2015;16:251. doi:10.1186/s13059-015-0824-9. PubMed PMID: 26578176; PubMed Central PMCID: PMCPMC4647450.

7. Zetsche B, Gootenberg JS, Abudayyeh OO, Slaymaker IM, Makarova KS, Essletzbichler P, et al. Cpf1 is a single RNA-guided endonuclease of a class 2 CRISPR-Cas system. Cell. 2015;163(3):759-71. doi:10.1016/j.cell.2015.09.038. PubMed PMID: 26422227; PubMed Central PMCID: PMCPMC4638220.

8. Gao L, Cox DBT, Yan WX, Manteiga JC, Schneider MW, Yamano T, et al. Engineered Cpf1 variants with altered PAM specificities. Nat Biotechnol. 2017;35(8):789-92. doi:10.1038/nbt.3900. PubMed PMID: 28581492; PubMed Central PMCID: PMCPMC5548640.

9. Ma S, Liu Y, Liu Y, Chang J, Zhang T, Wang X, et al. An integrated CRISPR Bombyx mori genome editing system with improved efficiency and expanded target sites. Insect Biochem Mol Biol. 2017;83:13-20. doi:10.1016/j.ibmb.2017.02.003. PubMed PMID: 28189747.

10. Isobe R, Kojima K, Matsuyama T, Quan GX, Kanda T, Tamura T, et al. Use of RNAi technology to confer enhanced resistance to BmNPV on transgenic silkworms. Arch Virol. 2004;149(10):1931-40. doi:10.1007/s00705-004-0349-0. PubMed PMID: 15669105.

11. Dong Z, Dong F, Yu X, Huang L, Jiang Y, Hu Z, et al. Excision of Nucleopolyhedrovirus Form Transgenic Silkworm Using the CRISPR/Cas9 System. Front Microbiol. 2018;9:209. doi:10.3389/fmicb.2018.00209. PubMed PMID: 29503634; PubMed Central PMCID: PMCPMC5820291.

12. Subbaiah EV, Royer C, Kanginakudru S, Satyavathi VV, Babu AS, Sivaprasad V, et al. Engineering silkworms for resistance to baculovirus through multigene RNA interference. Genetics. 2013;193(1):63-75. doi:10.1534/genetics.112.144402. PubMed PMID: 23105011; PubMed Central PMCID: PMCPMC3527255.

13. Dong Z, Huang L, Dong F, Hu Z, Qin Q, Long J, et al. Establishment of a baculovirus-inducible CRISPR/Cas9 system for antiviral research in transgenic silkworms. Appl Microbiol Biotechnol. 2018;102(21):9255-65. doi:10.1007/s00253-018-9295-8. PubMed PMID: 30151606.

14. Dong ZQ, Chen TT, Zhang J, Hu N, Cao MY, Dong FF, et al. Establishment of a highly efficient virus-inducible CRISPR/Cas9 system in insect cells. Antiviral Res. 2016;130:50-7. doi:10.1016/j.antiviral.2016.03.009. PubMed PMID: 26979473.

15. Chen S, Hou C, Bi H, Wang Y, Xu J, Li M, et al. Transgenic Clustered Regularly Interspaced Short Palindromic Repeat/Cas9-Mediated Viral Gene Targeting for Antiviral Therapy of Bombyx mori Nucleopolyhedrovirus. J Virol. 2017;91(8). doi:10.1128/JVI.02465-16. PubMed PMID: 28122981; PubMed Central PMCID: PMCPMC5375672.

16. Dong Z, Qin Q, Hu Z, Chen P, Huang L, Zhang X, et al. Construction of a One-Vector Multiplex CRISPR/Cas9 Editing System to Inhibit Nucleopolyhedrovirus Replication in Silkworms. Virol Sin. 2019;34(4):444-53. doi:10.1007/s12250-019-00121-4. PubMed PMID: 31218589; PubMed Central PMCID: PMCPMC6687805.

17. Dong Z, Hu Z, Qin Q, Dong F, Huang L, Long J, et al. CRISPR/Cas9-mediated disruption of the immediate early-0 and 2 as a therapeutic approach to Bombyx mori nucleopolyhedrovirus in transgenic silkworm. Insect Mol Biol. 2019;28(1):112-22. doi:10.1111/imb.12529. PubMed PMID: 30120848.

18. Dong D, Ren K, Qiu X, Zheng J, Guo M, Guan X, et al. The crystal structure of Cpf1 in complex with CRISPR RNA. Nature. 2016;532(7600):522-6. doi:10.1038/nature17944. PubMed PMID: 27096363.

19. Fonfara I, Richter H, Bratovic M, Le Rhun A, Charpentier E. The CRISPR-associated DNA-cleaving enzyme Cpf1 also processes precursor CRISPR RNA. Nature. 2016;532(7600):517-21. doi:10.1038/nature17945. PubMed PMID: 27096362.

20. Nakade S, Yamamoto T, Sakuma T. Cas9, Cpf1 and C2c1/2/3-What’s next? Bioengineered. 2017;8(3):265-73. doi:10.1080/21655979.2017.1282018. PubMed PMID: 28140746; PubMed Central PMCID: PMCPMC5470521.

21. Mahfouz MM. Genome editing: The efficient tool CRISPR-Cpf1. Nat Plants. 2017;3:17028. doi:10.1038/nplants.2017.28. PubMed PMID: 28260792.

22. Perez-Pinera P, Ousterout DG, Gersbach CA. Advances in targeted genome editing. Curr Opin Chem Biol. 2012;16(3-4):268-77. doi:10.1016/j.cbpa.2012.06.007. PubMed PMID: 22819644; PubMed Central PMCID: PMCPMC3424393.

23. Morisaka H, Yoshimi K, Okuzaki Y, Gee P, Kunihiro Y, Sonpho E, et al. CRISPR-Cas3 induces broad and unidirectional genome editing in human cells. Nat Commun. 2019;10(1):5302. doi:10.1038/s41467-019-13226-x. PubMed PMID: 31811138.

24. Moon SB, Kim DY, Ko JH, Kim YS. Recent advances in the CRISPR genome editing tool set. Exp Mol Med. 2019;51(11):130. doi:10.1038/s12276-019-0339-7. PubMed PMID: 31685795; PubMed Central PMCID: PMCPMC6828703.

25. Shen W, Zhang J, Geng B, Qiu M, Hu M, Yang Q, et al. Establishment and application of a CRISPR-Cas12a assisted genome-editing system in Zymomonas mobilis. Microb Cell Fact. 2019;18(1):162. doi:10.1186/s12934-019-1219-5. PubMed PMID: 31581942; PubMed Central PMCID: PMCPMC6777028.

26. Matsoukas IG. Commentary: RNA editing with CRISPR-Cas13. Front Genet. 2018;9:134. doi:10.3389/fgene.2018.00134. PubMed PMID: 29722368; PubMed Central PMCID: PMCPMC5915479.

27. Schindele P, Wolter F, Puchta H. Transforming plant biology and breeding with CRISPR/Cas9, Cas12 and Cas13. FEBS Lett. 2018;592(12):1954-67. doi:10.1002/1873-3468.13073. PubMed PMID: 29710373.

28. Savage DF. Cas14: Big Advances from Small CRISPR Proteins. Biochemistry. 2019;58(8):1024-5. doi:10.1021/acs.biochem.9b00035. PubMed PMID: 30740978; PubMed Central PMCID: PMCPMC6924505.

29. Kleinstiver BP, Tsai SQ, Prew MS, Nguyen NT, Welch MM, Lopez JM, et al. Genome-wide specificities of CRISPR-Cas Cpf1 nucleases in human cells. Nat Biotechnol. 2016;34(8):869-74. doi:10.1038/nbt.3620. PubMed PMID: 27347757; PubMed Central PMCID: PMCPMC4980201.

30. Wang F, Xu H, Yuan L, Ma S, Wang Y, Duan X, et al. An optimized sericin-1 expression system for mass-producing recombinant proteins in the middle silk glands of transgenic silkworms. Transgenic Res. 2013;22(5):925-38. doi:10.1007/s11248-013-9695-6. PubMed PMID: 23435751.

31. Smith DB, Johnson KS. Single-Step Purification of Polypeptides Expressed in Escherichia-Coli as Fusions with Glutathione S-Transferase. Gene. 1988;67(1):31-40. doi:Doi 10.1016/0378-1119(88)90005-4. PubMed PMID: WOS:A1988P255900004.

32. Pan MH, Cai XJ, Liu M, Lv J, Tang H, Tan J, et al. Establishment and characterization of an ovarian cell line of the silkworm, Bombyx mori. Tissue Cell. 2010;42(1):42-6. doi:10.1016/j.tice.2009.07.002. PubMed PMID: 19665160.

33. Zhang J, Chen XM, Zhang CD, He Q, Dong ZQ, Cao MY, et al. Differential susceptibilities to BmNPV infection of two cell lines derived from the same silkworm ovarian tissues. PLoS One. 2014;9(9):e105986. doi:10.1371/journal.pone.0105986. PubMed PMID: 25221982; PubMed Central PMCID: PMCPMC4164443.

34. Dong ZQ, Zhang J, Chen XM, He Q, Cao MY, Wang L, et al. Bombyx mori nucleopolyhedrovirus ORF79 is a per os infectivity factor associated with the PIF complex. Virus Res. 2014;184:62-70. doi:10.1016/j.virusres.2014.02.009. PubMed PMID: 24583368.

35. Naito Y, Hino K, Bono H, Ui-Tei K. CRISPRdirect: software for designing CRISPR/Cas guide RNA with reduced off-target sites. Bioinformatics. 2015;31(7):1120-3. doi:10.1093/bioinformatics/btu743. PubMed PMID: 25414360; PubMed Central PMCID: PMCPMC4382898.

36. Stemmer M, Thumberger T, Del Sol Keyer M, Wittbrodt J, Mateo JL. CCTop: An Intuitive, Flexible and Reliable CRISPR/Cas9 Target Prediction Tool. PLoS One. 2015;10(4):e0124633. doi:10.1371/journal.pone.0124633. PubMed PMID: 25909470; PubMed Central PMCID: PMCPMC4409221.

